# Odor tracking in flying *Drosophila* requires visual reafference and compass neurons

**DOI:** 10.64898/2026.03.13.711693

**Authors:** John Paul Currea, Ava E Bignell, Isabelle Rieke-Wey, Daniela Limbania, Brian J Duistermars, Sara M Wasserman, Mark A Frye

## Abstract

Flying *Drosophila* critically depend on high-contrast visual surroundings to localize odor sources in still air, yet the neural mechanisms of visual integration for active odor tracking are unknown. We demonstrate that E-PG neurons—head direction cells in the central complex—work in concert with self-generated visual motion signals to maintain a stable heading metric during olfactory navigation in flight. Using a magnetic tether system and a digital “visual clamp”, we show that removing the visual feedback generated by a fly’s own turns (reafference) causes the animal to lose its heading within an odor plume. Thus, olfactory and mechanosensory signals alone are insufficient for plume stabilization. E-PG neurons have been shown to store visual changes in heading during flight. Genetically hyperpolarizing E-PG neurons significantly compromised the flies’ ability to both acquire and maintain heading toward a food odor. Notably, silencing these neurons did not disrupt basic visual reflexes, such as optomotor gaze stabilization or object tracking, indicating a specific role in odor-directed visual navigation rather than basic visual flight control. While odor was found to modulate the frequency and amplitude of turns independently, E-PG neurons are essential for directing the orientation of corrective saccades toward the plume center. These results establish that visual reafference engages the internal visual compass to sustain a spatial working memory of heading changes between saccades, allowing flies to maintain a straight course and navigate effectively toward an invisible odor source in flight.

**Significance Statement:** A crucial brain function that remains largely mysterious is building robust working memory by integrating stimuli across sensory modalities. For flying *Drosophila*, visual feedback is required to localize an odor source. Recently, a group of cells residing in the navigation center of the fly brain (E-PG neurons) have been shown to function like an internal compass, providing a dynamic working memory of heading changes akin to mammalian head-direction cells. Here, we show that self-generated visual motion signals work with E-PG compass neurons so that the direction of exploratory turns can be stored and recalled to maintain the animal’s heading within an odor plume during flight.

## Introduction

Flying insects, such as *Drosophila*, require olfactory cues to locate food and reproductive sites, sometimes over vast distances (1). Odor plumes in nature can be intermittent and filamentous, so odor detection alone does not reliably indicate the source location because the structure of the odor plume is determined by wind conditions, which may rapidly shift direction or be entirely still (2). In wind, a flying fly tends to ‘cast and surge’, making cross-wind turns if the odor signal is lost (cast), and flying upwind while the odor is detectable (surge) (3, 4). In fact, because wind can determine the direction of an odor signal, odor activates central brain circuits that encode the relative wind direction for this purpose (5). The absence of wind poses a challenge to olfactory navigation that can be solved by measuring the static odor gradient and orienting up it (chemotaxis), or by changing the rate of turns as the odor intensity changes (chemokinesis). Walking flies use both strategies (6, 7). In flight, rapid turns—termed “saccades” for their functional similarity to gaze stabilizing eye movements—are more frequent and larger in still air than wind (8), emphasizing the importance of this motor behavior for odor tracking. Flying through still air, a food odor carried by advection tends to provoke a “sink and circle” behavior composed of decreased altitude (sink), coupled with bouts of unidirectional saccades (circle) (8, 9). Saccade direction is influenced by the static odor gradient across the two antennae, generating differential neural activation across the antennal lobes (7, 10).

Olfaction is essential in many ways and yet is insufficient on its own to enable odor localization during flight. Rather, *Drosophila* have a peculiar dependence upon vision to localize an odor source. In free-flight within a visually featureless, low contrast 1-meter arena, flies fail to target the advecting odor source in still air, yet they have no trouble approaching the source when the arena walls are covered with a high contrast pattern of vertical edges (9). Follow-up work tested hungry flies tethered on a magnetic pivot, free to steer in the horizontal (yaw) plane during flight. A slow, effectively windless, plume of food odor elicits small amplitude low frequency saccades directed back and forth across the plume. However, if the high contrast visual background is switched to a uniform grayscale, the animals no longer balance saccade direction to maintain their heading toward the plume, instead randomly exploring the circular arena as if there were no odor present (11, 12). The same visual dependence is observed with thirsty flies seeking water vapor in flight (13). Computational models indicate that subtle modulation of visual control parameters is sufficient to enhance odor localization in flight in still air (9, 14), yet the functional role and neural mechanisms of visually-guided plume tracking are unknown.

In flies and other insects, allocentric heading changes are encoded and stored by compass neurons that project from a toroidal neuropile called the Ellipsoid Body to the Protocerebral Bridge and Gall (15), thereby named “E-PG” neurons. During flight under head-fixed virtual reality conditions, a revolving bump of E-PG neural activity dynamically tracks the displacement of the visual scene as a fly alters its wing kinematics to steer back and forth in yaw, much like the needle on a compass except with each individual calibrated to its own “north” (16, 17). For a head-fixed fly, the time course and magnitude of bump displacement faithfully follows its voluntary course changes in flight. Most notably, segments of straight flight interspersed with spontaneous turns are tracked by phase-shifts in bump position, thereby providing a stable metric of heading direction even for turns separated by several seconds or more (17). We reasoned that high contrast stationary surroundings provide visual feedback about self-induced displacements and that these reafferent visual signals engage compass neurons to record heading changes, directing saccades into an invisible plume in still air. Here we show that removing visual reafference—visual input generated by the fly’s volitional turns—causes flies to lose the plume, and that hungry flies with silenced E-PG neurons are unable to locate an odor plume nor follow a food odor plume they have already encountered.

## Results

### Hungry magnetically tethered flies maintain their heading into a food odor plume

Flies were presented with a visual panorama projected from a digital projector, reflected from first-surface mirrors, to form a geometrically perspective corrected image that subtended 360° in azimuth and 70° in elevation (Figure 1A). A small nozzle supplied saturated vapor at 14ml/min flow rate, drawn away from beneath the fly with gentle suction. The fly is tethered to a steel pin and suspended within a frictionless magnetic pivot that allows free rotation in the yaw plane (Figure 1B). A photoionization detector was used to sample the plume gradient inside the flight arena, which indicates the highest concentration occurs near the odorant nozzle, but that trace odor stimuli occur across the azimuth (Figure 1C). The fly’s heading is measured using a high speed infrared video camera from below (not shown). To eliminate self-generated image motion during free-yaw turns, the yaw position of the fly is subtracted from the orientation of the stimulus in the next frame. In effect, this iteratively revolves the visual display in register with the fly’s yaw turns, removing the expected visual reafference but leaving proprioceptive sense of body rotation intact (Figure 1D). We performed an offline validation of the spatial and temporal precision of the clamp by using DeepLabCut (18, 19) to track body angle from the raw video footage. We then compare the offline tracking results to the original online heading measurement and resultant image motion dynamics (Figure 1E). Hardware projection error amounted to half of the acceptance angle of a single photoreceptor and less than one fifth of the impulse response time of an optomotor reflex (see Methods). Visual stimuli were composed of either a symmetric 30° wavelength squarewave grating or asymmetric natural panoramic image (Figure 1F). Two of the experiments below were performed using a 96×16 pixel cylindrical array of blue LEDs with 3.75 degree pixels equipped with identical magnetic tether and odor plume apparatus (Figure 1G)(13).

**Figure 1.**
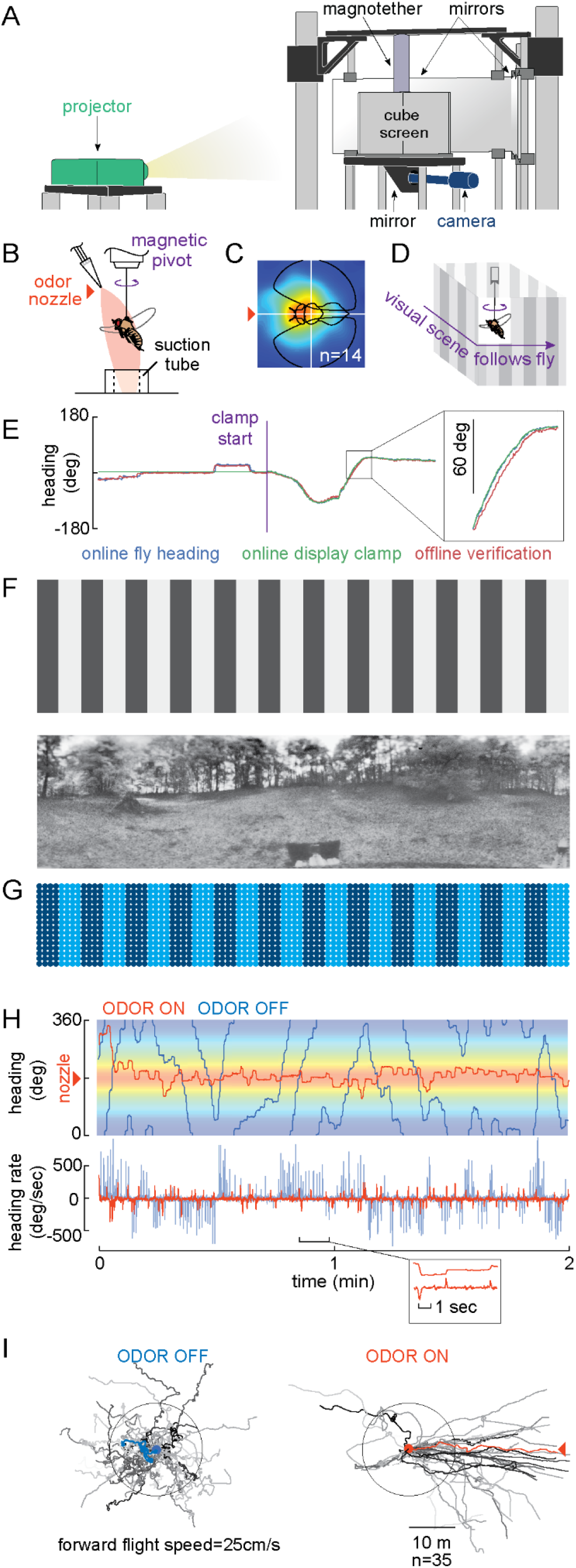
Flies actively center a food odor plume on a free-yaw visual flight simulator. (A) Digital projection onto mirrors forms a perspective corrected display on four sides of an acrylic cube lined with a diffuser. The display image was 1280 x 800 pixels, each subtending 0.4 degrees. (B) Saturated odor vapor (apple cider vinegar, ACV) is delivered from a nozzle 3 mm from the fly’s head and drawn away with suction. A magnetic pivot allows the fly or revolve 360°. (C) The spatial odor gradient were collected from a 12×12 grid using a photoionization detector. The grid was sampled 14 times and values were averaged at each point and plotted in pseudocolor. The fly silhouette is drawn to scale and the center of rotation indicated with the white crosshair. (D) The fly’s body angle is tracked in real time with infrared video and used to yoke the rotation of the visual display so that the visual scene remains stable relative to the fly’s body axis as the fly freely steers in yaw. (E) Time series plot of example fly heading, online display dynamics of the visual clamp, and offline fly heading tracked with Deep Lab Cut to validate online stimulus dynamics. Inset shows a zoomed-in segment indicative of the difference between fly heading and display movement, which was on average 2.6 degrees for 50% of all trials. (F) Two visual stimuli were programmed with the digital projection system, a 30 degree wavelength square grating and a natural scene. (G) Data in Figure 1H and Figure 4 were collected on a legacy system based on a cylindrical array of 24×96 blue pixels, subtending 3.75 degrees, drawn to scale. (H) Data from a single fly for a 2 minute trial in ACV and using the blue LED display. (I) For 35 flies tested as in G, simulated flight paths were reconstructed assuming a fixed forward flight speed. Each trajectory started from the same initial orientation. Circle indicates 10m radius.

To highlight the nature of plume tracking behavior, a plume of either saturated water vapor (odor OFF) or saturated apple cider vinegar (ACV, odor ON) was presented continuously for 2-minute trials. The visual panorama consisted of a stationary 30° grating on the LED display (Figure 1G). In the absence of ACV, an exemplar animal explores the 360° arena with rapid body saccades, characterized by spikes in angular velocity, which punctuate periods of stationary, fixed heading. When the odor plume is switched on, the saccades become smaller, less frequent, and are oriented to maintain the flight heading centered near the nozzle (Figure 1H). For these tethered flies that can change only their azimuthal heading, we simulated 2D flight trajectories *v* from yaw body angle θ assuming a moderate flight speed *r* =25cm/s (3, 4). Flight vectors were calculated for each time point with the following formula:

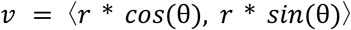

We tested 35 flies before and after odor exposure (Figure 1I). In the absence of odor, the simulated flight trajectories are short, curved or circular, and project in all directions from the starting point. By contrast, after being presented with the odor plume, flies balance the distribution of right and left saccades in order to maintain their heading toward the odor nozzle for 2 minutes, equivalent to roughly 30 meters at nominal flight speed of 25cm/sec (Figure 1I).

### Free-yaw flies use visual reafference to maintain a constant heading within a plume

Prior work using the free-yaw magnetic tether system has demonstrated the importance of visual reafference in odor tracking behavior. For flies that begin test trials within the odor plume, switching from a high contrast grating to an equiluminant grayscale background significantly reduces their ability to maintain their heading in the plume, and this visual dependence occurs both to track attractive apple cider vinegar and anti-track aversive benzaldehyde (11–13, 20). These results suggest that visual feedback generated during self-motion is somehow being stored, sometimes for several seconds, in order to direct saccades back into the plume. Therefore, removal of visual contrast eliminates the directional memory trace required to maintain heading. Alternatively, the spatially uniform visual panorama could trigger kinesis-like changes in saccade rate or amplitude that are independent of the control of heading direction.

To test these alternatives, we removed visual reafference normally generated during turns by digitally yoking or clamping yaw rotation of the visual display to the fly’s body axis in free-yaw conditions (Figure 1E). Thus, during a leftward saccade, the fly experiences normal changes in yaw inertia and odor intensity, but loses the expected rightward image displacement. We used the symmetrical striped grating background, which contains no spatial features that might impose a preferred heading.

Hungry flies (see Methods) were drawn to the ACV plume with bar oscillation before each trial. Each fly was tested both without the visual clamp and with the clamp starting at the 20 second mark (Figure 2A; from here onward, time series data are plotted vertically to facilitate visual inspection of left-right steering asymmetries across the midline odor plume). Visual inspection of the raw trajectories indicates that flies disperse from the midline plume when the clamp is activated (Figure 2A *right*), drifting when they would normally fixate. We calculated the probability of flight heading over the 360° visual azimuth for ten second epochs at the start (t=0-5 sec) and end (t=15-20 sec) of each trial (Figure 2B). The proportion of trajectories that centered the plume was reduced significantly by the clamp onset (Figure 2B *right*). However, we note the non-zero probability of transit near the plume even with the visual clamp engaged, suggesting some influence of visually-independent chemotaxis, at least over the 10 second time course of the test.

**Figure 2.**
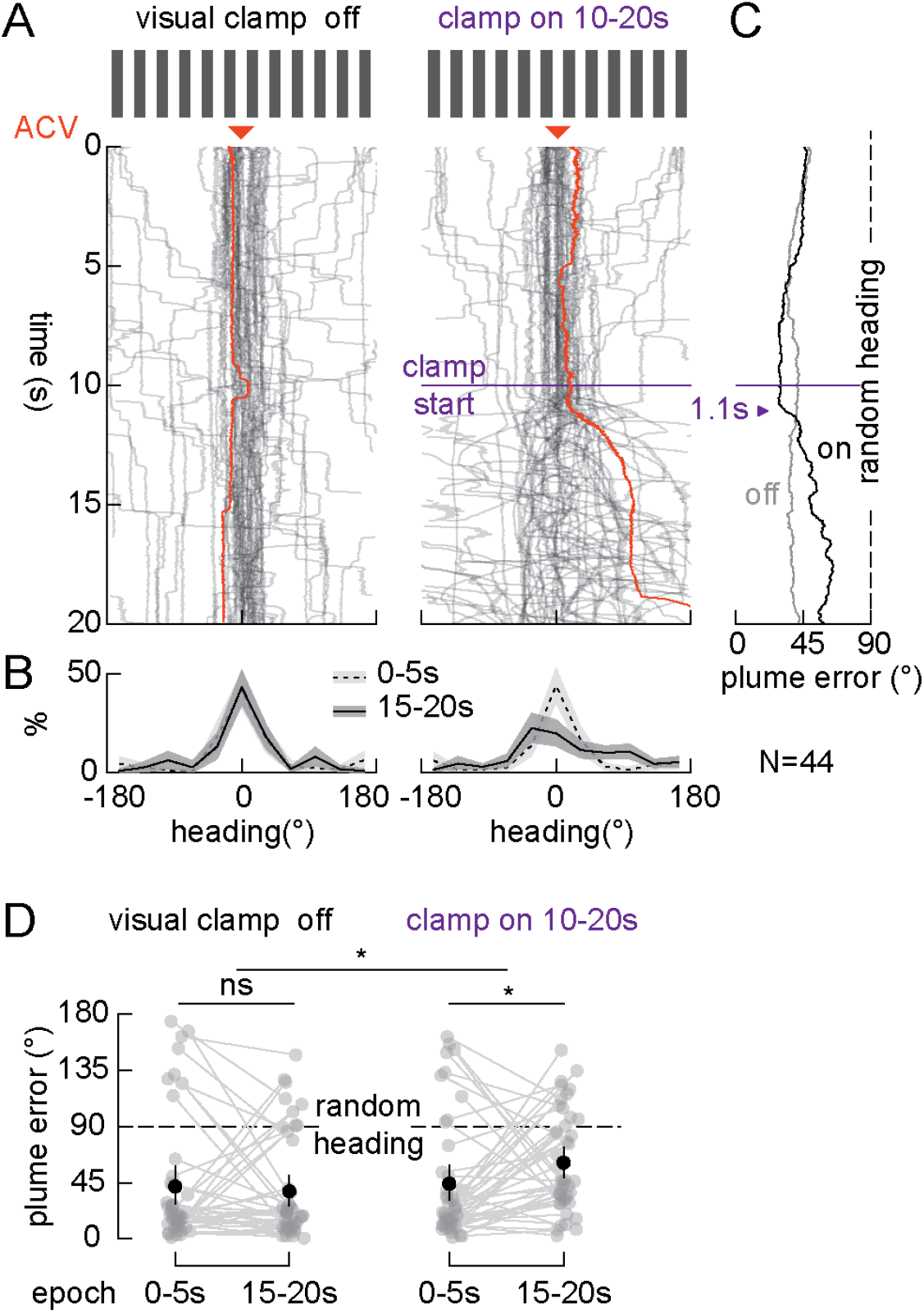
Removing visual reafference compromises odor tracking in flight. Flies were tested with a 30 degree symmetrical grating under continuous ACV delivery. (A) Raw flight trajectories for each fly in gray. One example trajectory is plotted in orange. Visual clamp remained off for the full trial (left) or was activated at the 10 second mark (right). Each fly was tested once in both conditions. Orange arrowhead indicates location of odor nozzle delivering ACV. (B) The distribution of heading across the population for two epochs: 0-10 seconds (dashed line) and 30-40 seconds (solid line). Shaded envelope indicates 84% confidence intervals, where non-overlapping indicate alpha<0.05. (C) The absolute angular distance from the plume averaged in time across the population. 90 degree plume error is equivalent to chance heading relative to the plume. (D) Average plume error across the test trial for each fly, within subjects comparisons across the first and last epoch of the trial. Error bars indicate 95% confidence intervals. Asterisks indicate level of significance of one-tailed bootstrap test (see Methods) *p<0.05.

To further quantify odor tracking capability, we computed the plume error as the angle between the fly’s heading and the odor nozzle. Plume error of 90° would be equivalent to chance heading relative to the plume (Figure 2C). Engaging the visual clamp at 10 s significantly increased plume error by 16° on average (P<0.05; Figure 2D), which led to an effect size of 20° more than the no-clamp test (P<0.05; Figure 2D). These results present direct evidence that when visual feedback experienced during closed-loop flight is disabled, olfactory and mechanosensory feedback signals alone are insufficient to stabilize heading into an odor plume. Rather, visual reafference generated specifically by self motion is crucial.

### Visual control of plume tracking requires compass neurons

We next sought to test whether the dependence on visual reafference for odor navigation required the use of E-PG compass neurons known to track visual heading changes, at least during body-fixed flight. We crossed a split-Gal4 line that selectively labels E-PG neurons (21) with UAS>Kir2.1, an inward rectifying potassium channel that hyperpolarizes neurons. F1 genetic controls carry a promoterless (empty) split-Gal4 at the same insertion driving UAS-Kir2.1, thereby obviating the need for parental controls with different genetic background.

The experimental design was similar to the visual clamp experiment above. However, since the angular domain of action of the E-PG activity bump can “jump” between symmetric stripes (22), we tested flies with both a periodic grating and a natural panorama. We compared the azimuthal distributions of flight heading during the initial 10 second epoch, when flies had been visually positioned near the plume, to the final 10 second epoch, when flies would be expected to be actively maintaining their heading. The empty>Kir control animals track the plume well within both the artificial grating and natural visual panoramas (Figure 3A), with no obvious differences in the trajectory of average plume error (Figure 3C). The fraction of flies maintaining their heading directly at the plume center was lower for the grating than for the natural scene (Figure 3B 30s→40s), see non-overlapping confidence intervals at 0°). On average, plume error increased roughly 15° over the course of the experiment, but this was not significant (P>.05) and there were no significant effects of switching between visual backgrounds for the genetic control flies (Figure 3D). The results indicate that both the grating and natural scene are effective at facilitating plume tracking, although centering performance is improved with the natural scene, at least close to the odor nozzle.

**Figure 3.**
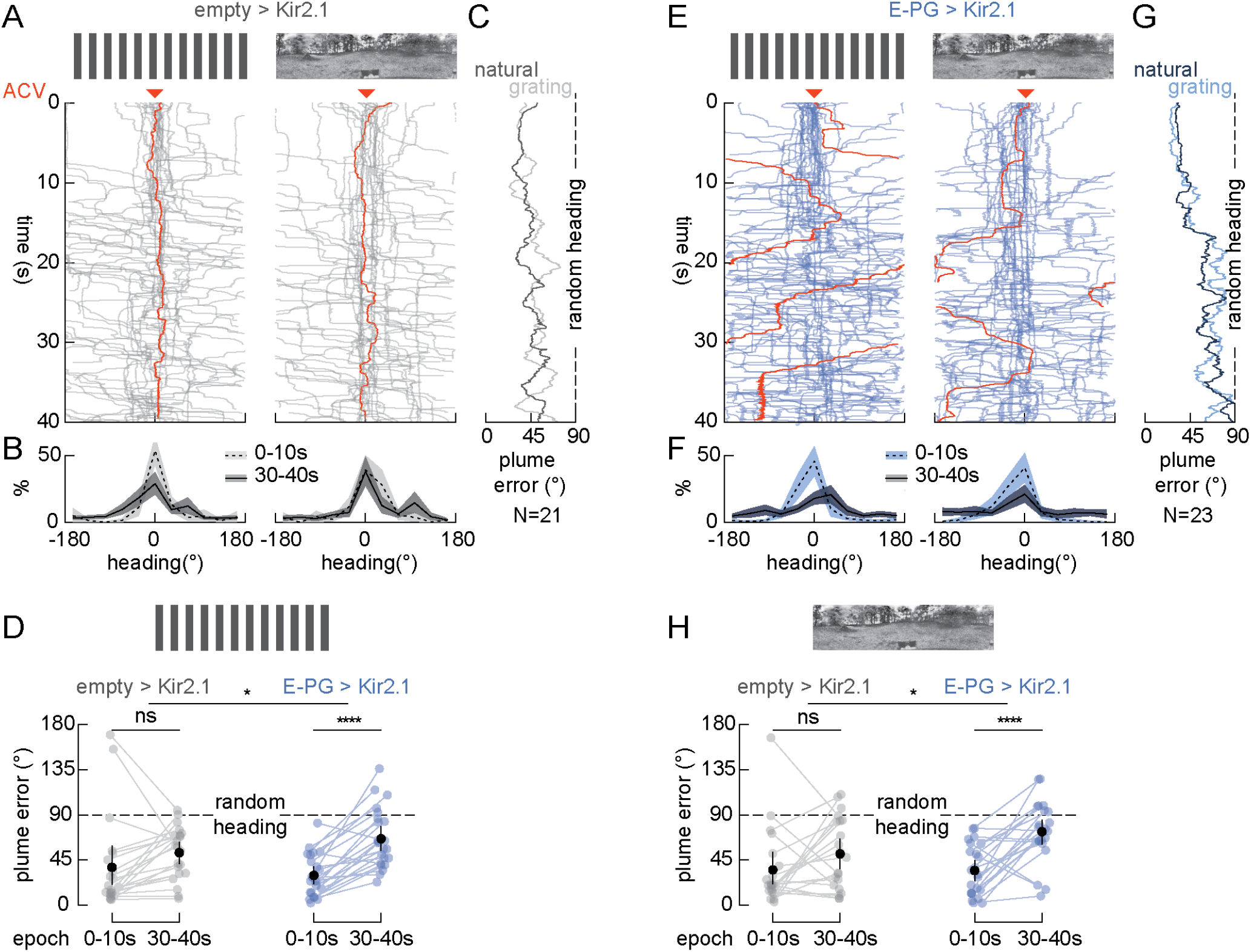
Compass neurons are required for flies to maintain normal plume centering behavior in flight. All panels are formatted the same as in Figure 2. (A) Empty-split-Gal4 controls are driving expression of UAS-Kir2.1; shorthand is empty>Kir2.1. Raw flight trajectories for each fly in gray. Flies were tested with both a 30 degree symmetrical grating (left) and natural scene (right). Orange arrowhead indicates location of odor nozzle delivering ACV. (B) Heading histograms with 84% confidence intervals, as in Figure 2. (C) Plume error averaged across the population as in Figure 2. (D) Plume error across the trial for each fly (dot), tested as in Figure 2. (E,F,G,H) Same plots as A,B,C,D but for E-PG-split-Gal4 driving expression of UAS-Kir2.1 shorthand E-PG>Kir2.1.

Silencing E-PG neurons with Kir2.1 strongly compromised flies’ ability to track the odor plume. Visual inspection of the raw trajectories suggests broad dispersion of flies’ heading away from the odor nozzle during the time course of the experiment (Figure 3E). This is evident in the heading histograms comparing the first and last 10-second epochs, which show that centering probability is reduced significantly between epochs for EPG>Kir flies (Figure 3F, notice that the confidence intervals don’t overlap around x=0°). Plume error accumulates throughout the trial approaching chance level (Figure 3G). On average, plume error increases 37° and 39° between the start to the end of the experiment for the grating and natural scene, respectively, conferring a high level of statistical significance (Figure 3H).

Silencing E-PG neurons significantly impairs, but does not eradicate, plume centering behavior, as evidenced by a significant number of trajectories centered at 0° (Figure 3F). Because flies were motivated to start close to the source of the plume, these experiments specifically tested whether they could maintain their heading in the food odor plume, not whether they could actively acquire it. To test for plume acquisition and simultaneously replicate the results, we performed the next experiment using different individual lab members, with a new genetic cohort, on a separate flight arena apparatus housed in a different lab room. This apparatus has been used in several other projects from the lab (10–12, 20), with identical olfactometer hardware, but is equipped with a 16×96 wrap-around dot matrix LED display that cannot present high resolution natural scenes. We used a high contrast 30 degree squarewave grating typical of other studies from our lab. All aspects of this experiment were similar to those in Figure 3, except that flies were not visually aligned into the plume.

In this dataset, flies were tested repeatedly allowing us to separate responses into two groups based on whether they started less than or greater than 90° from the nozzle. Control flies show such fast responses to the odor that we had to reduce the first epoch down from 0-5 s to 0-1s in order to properly capture the initial heading distributions. The empty>Kir flies that started within 90° of the nozzle remained centered at the plume with no significant change in heading distributions from the start and end of the trial, whereas those that started from the periphery were able to center it (Figure 4A,B), highlighted by the convergence of plume error for the two groups (Figure 4C). By contrast, the E-PG>Kir flies tended to maintain a high error throughout the trial after starting in the periphery and did not center it (Figure 4E,F,G). Quantifying plume error across the population confirmed that starting at the plume can bias the measurements. For flies starting near the plume, E-PG silenced flies indeed perform nearly as well as controls, with roughly similar increases in plume error over the course of the experiment (Figure 4D). However, for those flies that are challenged to acquire the plume, the empty>Kir flies radically reduce their plume error, significantly below chance, whereas E-PG silenced flies perform at chance level (Figure 4H). These results demonstrate that hyperpolarizing E-PG neurons compromises both plume acquisition and active centering. Together with the visual reafference results in Figure 2, these indicate that visual input to working memory maintained by the local activity bump within E-PG compass neurons is crucial for flies to navigate an odor plume in still air.

**Figure 4.**
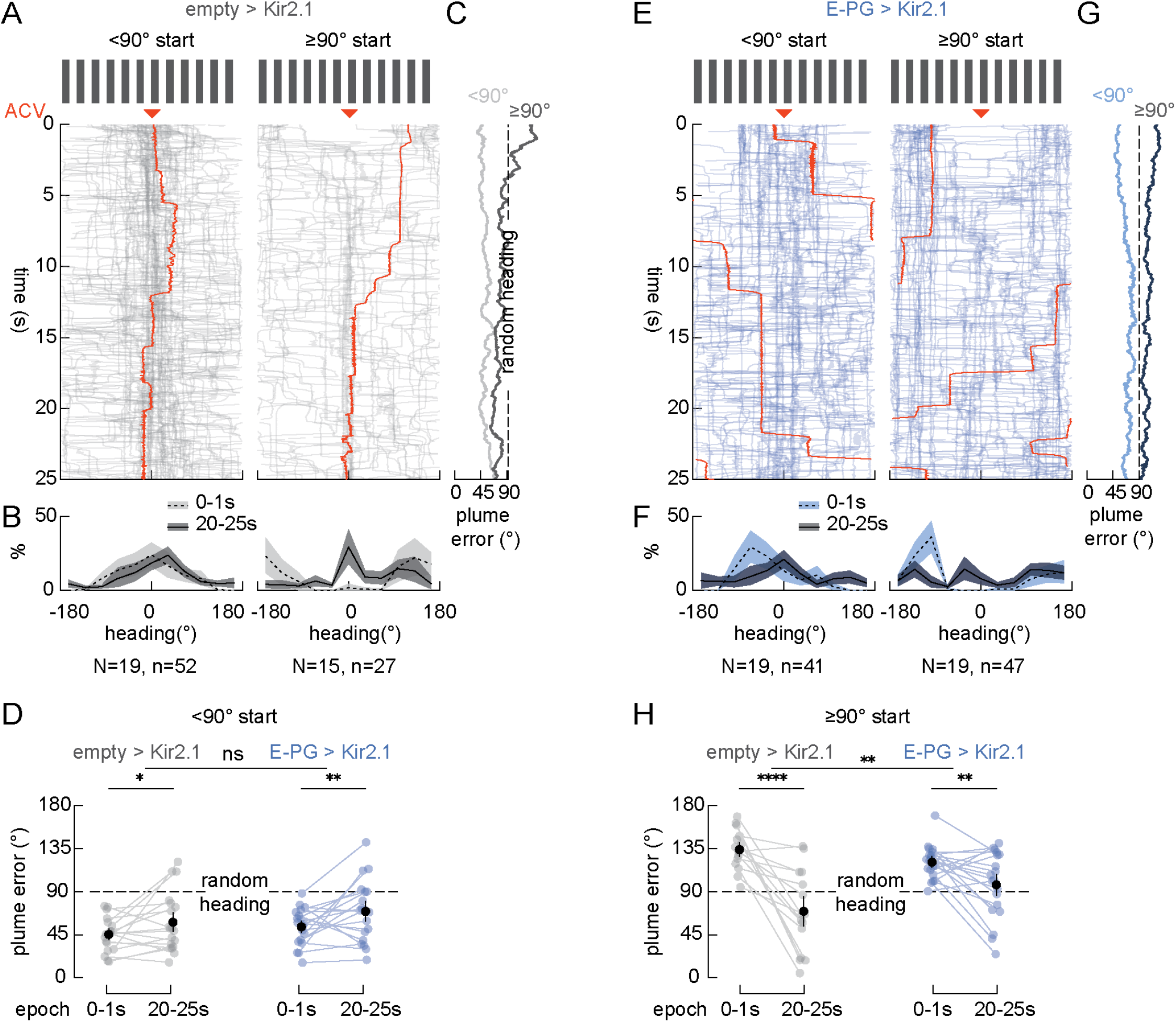
Compass neurons are required for flies to acquire the plume in flight. All panels are formatted the same as in Figures 2,3. (A) Flies were sorted into those that started trials near the odor nozzle (left) and those that started far from the nozzle (right). Orange arrowhead indicates location of odor nozzle delivering ACV. Example individual traces highlighted in orange. (B) Heading histograms divided for the first one second (dashed line) and last five seconds (solid line). (C) Average plume error as in Figures 2,3. (D) Plume error comparisons within subjects for flies starting trials near the odor nozzle. Error bars indicate 95% confidence intervals. Asterisks indicate level of significance of one-tailed bootstrap test (see Methods) *p<0.05, **p<0.01 (E,F,G) Same plots as A,B,C but for E-PG-split-Gal4 driving expression of UAS-Kir2.1. (H) Plume error comparisons within subjects for flies starting trials away from the odor nozzle. Difference in means was tested with bootstrap comparisons resampled 100,000 times. Error bars indicate 95% confidence intervals. **p<0.01, ****p<0.0001. Empty>Kir2.1 N=21 flies; E-PG>Kir2.1 N=22 flies.

### Silencing compass neurons does not disrupt visual flight control systems

Central complex circuitry has been shown to coordinate goal directed navigation behavior in insects as diverse as flies, locusts, butterflies, dung beetles, cockroaches, and bees (23). Thus, how can we be sure that disrupting the excitability of E-PG neurons does not compromise visual flight control in a manner that acts independently from goal directed navigation? To explore this possibility, we performed a set of control experiments on E-PG>Kir flies to assess wide-field optomotor gaze stabilization and small-field bar tracking - two robust visual behaviors that do not rely on spatial memory for flight control.

We first tested gaze stabilizing optomotor reflexes using a 30 degree grating rotating at constant speed. We find that E-PG>Kir flies perform as well as controls, tracking image motion with constant velocity smooth turns interspersed with occasional saccades that is typical of wild-type flies (24). The steady-state average responses of the two experimental groups overlap nearly perfectly (Figure 5A). Next, we assessed dynamic steering responses to brief impulses of wide-field image motion. Using an m-sequence to approximate white noise, we used reverse correlation to derive the optimal linear filter linking optic flow to changes in flight heading (25). The estimated filters for body heading changes in magnetically tethered free-yaw flies show onset delay of ~50 milliseconds and time-to-peak of ~150 milliseconds, which closely agrees with measurements from the wing steering kinematics of wild-type rigidly tethered flies; there were no significant differences between empty>Kir and E-PG silenced flies (Figure 5B).

**Figure 5.**
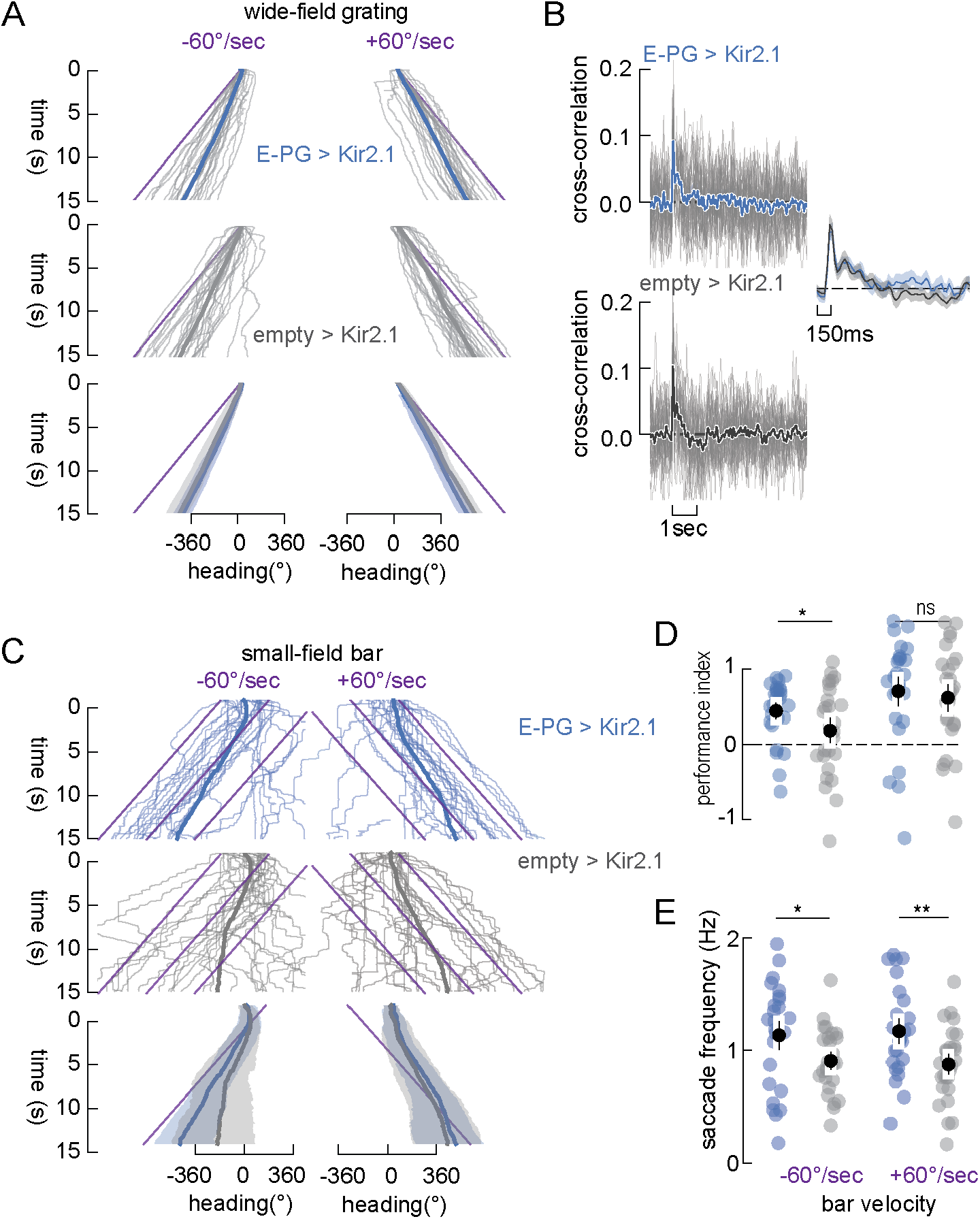
Basic visual flight control is not disrupted by silencing compass neurons. (A) Steady-state smooth optomotor stabilization test. A high resolution grating of 30 degree wavelength stripes (Figure 1F) was revolved at constant velocity. Individual fly responses plotted with thin lines. Average of the population in thick lines. Shaded envelope indicates 95% confidence intervals (bottom). Stimulus path indicated in purple lines. (B) Impulse response test. A 30 degree grating was moved along an m-sequence trajectory to simulate spectrally white oscillation frequencies. Individual fly responses plotted with thin lines. Average of the population in thick lines. Inset re-plots and overlays the average impulse responses for the two groups. Shaded envelope indicates 95% confidence intervals. (C) Bar tracking behavior test. A randomly textured 30 degree bar was moved across a similarly random textured stationary background at constant speed. Stimulus indicated in purple lines. Individual fly responses plotted with thin lines. Average of the population in thick lines. Shaded envelope indicates 95% confidence intervals (bottom). (D) Performance index equals the ratio of fly azimuthal displacement to bar displacement. For each fly (colored dot) and population means (black dot). (E) The rate of saccades during each trial, for each fly (colored dot) and population means (block dot). Difference in means was tested with bootstrap comparisons resampled 100,000 times. Error bars indicate 95% confidence intervals. *p<0.05, **p<0.01. Empty>Kir2.1 N=21 flies; E-PG>Kir2.1 N=22 flies.

Next, we tested bar tracking behavior. In free-flight and on a magnetic tether, *Drosophila melanogaster* does not smoothly track the trajectory of vertical edge objects resembling plant stalks, instead making orientation turns toward the object’s position while integrating the movement error to a threshold for executing a pursuit saccade (26, 27). Here, we presented a randomly textured bar superimposed upon a stationary similarly textured visual background revolving at constant speed. In response to this small-field motion stimulus, flies track the bar using saccadic changes in visual heading while fixating the visual surroundings in between saccades. Average change in flight heading did not differ between empty>Kir and E-PG>Kir flies (Figure 5C). We computed the performance index (28), the ratio of the fly’s angular displacement to that of the visual stimulus. PI=1 would be perfect bar following, whereas PI=-1 is steering in the opposite direction and PI=0 is no response to the bar. E-PG silenced flies did not show weakened PI values, and in fact were better at bar tracking at least in the negative direction (Figure 5D). Finally, we computed saccade rates during small-field bar motion trials. Here, E-PG silenced flies show significantly higher saccade rates, which we attribute to rigorous bar pursuit reflexes (Figure 5E). These control experiments indicate that silencing compass neurons has no obvious impact on flight stabilization reflexes or object approach preferences. Rather, compass silencing causes deficits specifically associated with goal directed navigation.

### The directional control of plume centering saccades requires compass neurons, but not their frequency or amplitude

The control variables that maintain flight heading in *Drosophila melanogaster* include reducing retinal slip to fly straight between rapid course changing saccades. Key saccade control parameters include amplitude, direction, and rate, which have been shown to be modulated by odor during flight (12, 13). We tested whether these variables are influenced by inactivating compass neurons. Figure 6 shows saccade peak velocity plotted at the azimuthal retinal position at which each saccade occurred. Note that we performed the analysis described below for each of the two visual treatments, the natural panorama and striped grating, and found that the results were statistically indistinguishable, so we pooled data across the two visual background treatments. Saccades are spatially localised in two quadrants of the plot indicating that when flight heading is oriented to the left of the plume (negative “saccade start position”) they tend to execute saccades to the right (positive “saccade peak velocity”), and *vice versa* (Figure 6A). To highlight the separation of saccade direction toward the plume, the data are re-plotted, zoomed-in on the central 180-degree azimuthal region (Figure 6B). The average saccade start position is separated by 36 degrees in azimuth, straddling the zero-position of the odor nozzle (Figure 6A,B empty>Kir2.1). Therefore, effector control flies make saccades directed toward the plume center. By contrast, the saccades evoked by E-PG blocked flies are not, on average, directed toward the plume. In fact, we observed a peculiar side-bias for the population of E-PG>Kir saccades, executed from the left side of the plume (Figure 6A,B E-PG>Kir2.1), which may be indicative of the disrupted directional control of saccades upon silencing E-PG neurons. We did not observe similar bias in other behavioral measurements.

**Figure 6.**
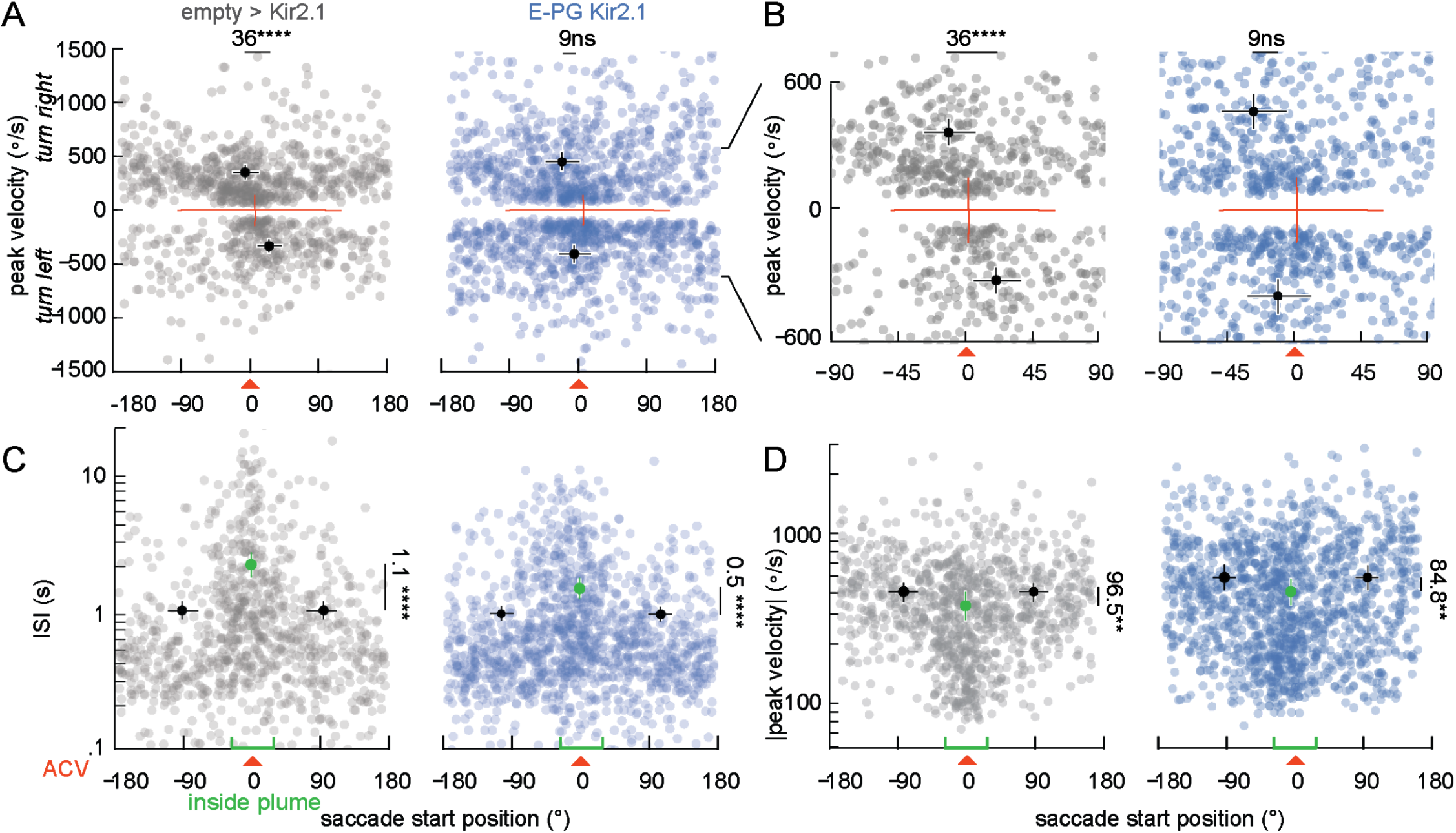
Compass neurons are required for normal saccade orientation toward the plume, but not for lower saccade rate within the plume. (A) For each saccade in the data set, the azimuthal angle at the start of each saccade is plotted against its signed peak angular velocity (i.e. positive values indicate rightward saccades). Orange cross hair indicates the zero axis intercepts. Orange arrowhead indicates location of odor nozzle delivering ACV. Dark dots indicate population means; error bars indicate 95% confidence intervals. Difference in means is indicated numerically at the top of each plot. (B) Data replotted from panel A but zoomed into a smaller range of axis values to highlight that flies oriented to the left of the plume made right directed saccades and *visa versa*. (C) For each saccade, the time interval to the prior saccades is plotted against the azimuthal position of fly heading at the start of the saccade. Mean values of saccades within +/-30 degrees of the odor nozzle (green X-axis marks, green symbols) were compared to those greater than 30-degrees (black symbols). Assuming bilateral symmetry, to compute peripheral means, data were reflected over the azimuth and pooled. Difference in means is indicated numerically at the right of each plot. (D) For each saccade, the absolute value of peak velocity is plotted against the azimuthal position of heading at the start of the saccade. Mean values are computed from two azimuthal groups as in panel C. In all panels, the difference in means is indicated with a solid line spanning the mean points. In all panels, difference in means was tested with bootstrap comparisons resampled 100,000 times. **p<0.01, ****p<0.0001. Empty>Kir2.1 n=997 saccades from N=21 flies; E-PG>Kir2.1 n=1418 saccades from N=22 flies.

We also compared inter-saccade intervals (ISI) across experimental conditions, because ISI has been shown to increase when a fly is inside a plume of any attractive odor so far tested (11). Increased ISI would help keep the fly oriented within the plume by pausing turns, a phenomenon observed in free flight within a linear wind tunnel (3). We parsed the ISI values into two groups: those located within +/-30 degrees of the plume, and those that occurred more than 30 degrees from the nozzle. There were no differences in the mean ISI (or peak velocity) on either side of the odor nozzle, so we reflected and pooled data to compute population averages. Mean ISI is significantly larger for saccades elicited near the plume center than in the periphery of the arena, and this change occurs for both control and E-PG silencing flies (Figure 6C), although it is significantly larger in control flies. We performed the same analysis on the distribution of peak angular speed—absolute value of peak angular velocity—for the two experimental treatments. The peak angular speed of saccades evoked when the fly is headed into the ACV plume is significantly smaller than for saccades executed in the periphery of the arena, and this effect persists for flies with silenced E-PG neurons (Figure 6D), although they were generally faster overall. Odor therefore reduces both saccade rate and amplitude independently from visual compass activity, indicating that some plume tracking control variables are under multisensory control (e.g. saccade direction, inter-saccade optomotor gain, odor activated object approach) whereas others are not (e.g. saccade interval, saccade amplitude).

## Discussion

There are several odor modulated visual behaviors that contribute to olfactory search and localization behavior during flight in flies. First, a fundamental visual reflex—the gaze stabilization by optomotor response that minimizes retinal slip to enhance visual detection, operates in flies as a linear transformation with high gain, and does not adapt to varying feedback conditions (25, 29, 30)—is strengthened by food odor in Drosophila (31). Computational modeling has demonstrated that even a small increase in odor modulated optomotor responses, iterated over many inter-saccade intervals, is sufficient to bring a fly closer to an odor source in flight (9, 14). This helps explain how flies maintain a forward heading when odor evokes an upwind surge (4, 32, 33). At least one wide-field directionally selective motion detecting neuron, residing within the visual pathway that is implicated in optomotor control, shows increased visual responses when visual stimuli are paired with food odor (34).

Odor encounters also engage robust visual fixation and centering of objects during flight. In free-flight, after encountering a food odor, flies approach a dark object on the arena floor that they ignore in clean air (4). A similar phenomenon is observed in tethered flight under visual closed-loop feedback conditions, in which food odor increases flies’ tendency to actively steer toward vertical bars and small spots (35, 36). It is noteworthy that the distribution of saccade direction with respect to visual features is qualitatively similar to the orientation functions toward olfactory features presented here (9, 37), highlighting shared motor control variables for flight navigation toward salient sensory features.

Visual reflexes in flies are fast - the full cascade of visual processing and musculoskeletal mechanics required to evoke a steering response by the wings in response to an impulse in image velocity occurs within 100 milliseconds (25). Olfactory motor responses are slower - a corrective turn into an odor plume during flight occurs with an onset delay of ~190 milliseconds and an a corrective turn back into the plume is delayed 450 milliseconds from loss of the odor signal (4). By contrast, the time intervals between saccades are at least ten times longer than these sensory reflexes. In free-flight, the distribution of inter-saccade interval peaks at roughly 700ms and tails off at 1.1 seconds (38). Our measurements of ISI inside the plume are longer still, ranging 300ms to 10s (Figure 1H inset, Figure 6C). After accounting for those neural processing delays, and notwithstanding some added sluggishness due to free-yaw tethering, that still leaves potentially multiple seconds during which a fly is maintaining a straight heading and must retain a working memory of heading change from the prior saccade. The compass has it covered. During flight under head-fixed virtual closed-loop flight conditions with a naturalistic visual scene, the activity bump of E-PG neurons maintains a faithful representation of an animal’s stationary heading for up to 10 seconds between turns (17).

We posit that visual reafference acts with E-PG signals to guide saccade direction to maintain heading up an odor plume. However, during voluntary saccades, pre-motor efference copy commands hyperpolarize directionally selective motion detecting neurons that coordinate gaze stabilizing head movements (39, 40). This mechanism is thought to enable flies to suppress the perception of visual motion during rapid body saccades (41) in much the same way that our own eye saccades are invisible to us. This suppressive effect on motion perception occurs only for voluntary changes in heading, not for reactions to external visual perturbations such as might occur after a gust of wind (42). By contrast, during head-fixed flight, the E-PG bump does not follow steering commands in the dark, and therefore it would appear that pre-motor commands alone are insufficient to spin the compass (43). It is unknown whether proprioceptive feedback from naturalistic body rotation during free-yaw saccades, perhaps originating from gyroscopic sense organs, might supply mechanosensory input that dynamically tunes the E-PG heading compass (43, 44).

An alternative explanation for our results is that visual reafference acts separately from non-visual mechanosensory input to the heading compass, and that perturbing each component evokes distinct neural disabilities that coincidentally compromise odor tracking behavior. However, one piece of evidence to suggest that visual reafference is indeed interacting with odor activated working memory of visual heading is that plume centering can persist long after the odor has been switched off, but does not persist after visual feedback is removed. Presenting a plume of ethyl butyrate, a liquid ester that smells like pineapple, evokes strong plume centering in our apparatus. However, unlike ACV or other food odors, switching off the ethyl butyrate plume does not immediately reduce plume centering behavior. Instead, flies continue to direct saccades toward the “phantom plume” for up to 25 seconds, unless visual feedback is removed by switching the visual display from a high contrast striped grating to zero-contrast grayscale (11). If proprioceptive signals were to be supplying the E-PG compass, then flies should be able to maintain heading in the “phantom plume” even within visually featureless surroundings.

For a walking fly, proper E-PG activity does not require visual input. Rather, even in darkness it encodes changes in walking direction, likely using proprioceptive feedback from the legs, although visual cues are weighted more than walking signals when these are in conflict (16). Regardless of how multisensory signals are balanced for mapping compass dynamics to body movement, the compass is crucial not only for odor navigation during flight, as we show here, but also when the fly is walking; recent results indicate that silencing E-PG neurons removes the capability of a fly to actively follow an odor plume while walking in the dark (6).

E-PG neurons are not the only components of the central complex that assist with olfactory navigation. A class of neurons intrinsic to the fan-shaped body (FSB) temporarily stores a fly’s heading direction upon encountering food odor presented within a wind stream, and this information is used to guide upwind navigation during walking (45). If FSB circuits could be activated by odor to store visual heading direction instead of wind direction, then this is precisely the sort of mechanism that could underlie visual navigation to an odor source in still air.

Our working model of visual-olfactory integration mechanisms for plume navigation consists of three components: 1) odor-modulated optomotor gain helps a fly maintain a straight course between saccades, 2) odor-amplified visual object saliency draws the animal’s attention and engages saccades toward otherwise neutral features, and 3) visual reafference-modulated saccade balancing, mediated by the EPG compass, maintains a stable flight heading in still air. Our results motivate additional research questions. 1) How does odor evoke a state change from exploration to heading-controlled tracking, in which the compass neurons balance saccade valence? 2) Does odor activate a saccadic search rhythm to probe the plume boundary? The corollary would be to simply fly straight in a plume. 3) How does the plasticity of heading circuitry facilitate olfactory navigation (46)? Exploring these questions likely involves examining the FSB circuits that (1) convey the optic flow that encodes flight travel direction independently from heading direction (47) and (2) are activated by odor to store wind direction information and mediate persistent upwind walking (5).

## Materials and Methods

### Fly stocks

Fly lines of adult female *Drosophila melanogaster*, 3–6 days posteclosion.

- *D. melanogaster: SS00096-SplitGAL4 (P{w[+mc]=GMR19G02-pBPp65ADZpUw} in attP40*
- *D. melanogaster: Empty-SplitGAL4 (P{w[+mc]=BP-p65ADZp} in attP40} and P{w[+mc]=BP-ZpG4DBD} in attP2)*
- *D. melanogaster: +;+;UAS-eGFP-Kir2*.*1 (pJFRC49-10XUAS-IVS-eGFPKir2*.*1 in attP2)*
- *D. melanogaster wild-type strain originating from a wild caught female in 2008*

### Flight experiments

Each fly was removed from food and water for at least one hour before testing. Flies were tethered as described previously(48, 49). Each fly was placed in the flight arena, and tested with a 30 degree grating revolving at constant speed in each direction. Visual inspection confirmed whether the fly could freely rotate and smoothly follow the stimulus at least one full revolution. At the start of each experimental trial, a 15 degree white vertical bar projected onto a solid black background was oscillated in front of the nozzle for five seconds to orient the fly toward the plume. Random block statistical design was used, and stimulus variables were presented within subjects for each experiment.

### Data, materials and software availability

All analyses were performed in Python, using a combination of established APIs (numpy, scipy, matplotlib, and statsmodels). Upon publication, data and plotting routines will be deposited in Open Science Framework (OSF).

### Magnocube

Recent work has demonstrated that coordinated photoreceptor mechanical contraction and intraretinal movements impart significantly higher spatial acuity than previously thought in Drosophila (42, 50). We adopted the high resolution projector display and graphics library of Cabrera and Theobald (51) and this system has been described previously (48). A digital projector and first-surface mirrors projects an image into the vertical sides of a 4’’x4’’x4’’ perspex cube lined with gray rear projection material. The projector (TI DLP LightCrafter 4500 EVM) produced frames 1280 x 800 pixels each subtending 0.4 degrees at 120 Hz. The magnetic tether was set up as described above, positioning the fly at the center of the cube. From this position, the display subtends the fly’s visual field by 360 horizontally and 70–90 vertically, decreasing from the center of each panel to the corners. This discrepancy is accounted for by programmatically restricting the vertical subtended angle to 70 degrees all around the azimuth and correcting for perspective. 12 concentric IR LEDs illuminate the magnet behind the fly while the fly sits in their shadow, cast by the bottom magnet, so the lights don’t illuminate the fly directly. Instead, we use the shadow of the fly, which contrasts greatly against the diffuse and reflected IR light. The brightness of the LEDs and camera exposure times were customized to minimize the influence of motion blur and shot noise.

### Realtime Tracking

To track the flies in realtime, we added custom Python code to the Holocube library. Holocube uses pyglet (https://pyglet.org/) and OpenGL (https://www.opengl.org/) to render perspective-corrected viewports of 3D graphics and a range of psychophysical stimuli. Holocube runs on an event loop at about 60Hz in our system and our code adds a function to each loop during the acquisition phase of each stimulus. When a new fly is added, they are shown a rotating dot field that flips direction every 8 seconds to both assess the subject’s responsiveness and tune the apparatus. As this runs, the experimenter adjusts the position of the camera using micromanipulators until the center of rotation aligns as close as possible to the center of the camera. The tracking algorithm relies on thresholding each frame to separate the fly (except for their wings) from the background. A graphical user interface (GUI) allows the experimenter to adjust this threshold while watching the fly respond to the widefield rotation in realtime. This GUI also allows the user to adjust the size of two concentric rings at the center of the frame, which are used for measuring the fly’s heading. Due to the way the flies are tethered, their abdomen sticks out a bit further from the center of rotation than their head. So, the outer ring first determines the tail angle based on the circular mean of the angles corresponding to the thresholded values. Then, after omitting the tail angle +/-90 degrees, the inner ring determines the head angle by the same process. This is repeated every loop and allows very rapid heading calculations (up to at least 480 Hz) because it converts a typically 2D process into a mostly 1D one.

However, unintentional translations run the risk of being misinterpreted as rotations, muddying the results, and side asymmetries due to off-center tethering run the risk of shifting the measured angle away from the true angle by some offset. So, we built an offline DeepLabCut (18, 19) model to validate the results of this online tracking algorithm. We found that the mean difference between the realtime and validated offline tracking across trials amounted to 0.04 degrees (5.58 standard deviation). The 50th percentile of the distribution of mean error was 2.6 degrees, and the average frame interval was 17 milliseconds (2.26 standard deviation). Thus, the realtime tracking operated faster than the onset delay of the visuo-motor impulse response in flight (25), and the spatial error amounted to less than the acceptance angle of a single ommatidium(52). All of the code for is freely available on github (https://github.com/jpcurrea/magnocube).

### Visual Clamp

We modified our digital projection system to deliver high resolution open-loop “clamped” visual stimuli to the free-yaw fly to subtract expected visual reafference (53). This system has been described previously (48). Briefly, using the same realtime tracking described above, this system subtracts its generated motion by the next frame. To do this for each event loop, the system sets the angular coordinate of the background, specifically about the vertical z axis, equal to the heading measured from the most recently captured image from the camera. Because the realtime tracking and display system generated negligible spatiotemporal error (mentioned above and in Figure 1E), setting the background coordinate system equal to the fly’s frame of reference virtually opens the visual feedback loop, effectively eliminating visual reafference.

It is worth noting that the display’s center of rotation matches the fly’s center of rotation at the tether point, not the center of rotation of the eye. Thus, when the display is stationary and the fly rotates, an object appearing near visual midline would dilate and contract slightly as the fly rotates towards and away. The radius from the fly’s head to the thorax is no more than 0.5 mm and the radius of the arena is 50.8mm. The greatest ratio of the subtended angles (maximum/minimum) is 1.02, so there’s a 2% distortion in the size of frontal versus rear objects. For the stripes of the 30 degree grating, this means true perceived stripe widths ranging from 14.85 to 15.15 degrees, a range of 0.3 degs, between stripes in front and behind the fly, respectively. That is an order of magnitude less than the acceptance angle of a single photoreceptor and two orders of magnitude less than the nominal pattern wavelength. These distortions are sufficiently small to be negligible. Furthermore, with the clamp activated, because the center of fly rotation is the center of the arena, then the minimally distorted size of whatever is in front of the fly maintains the same angle as the fly rotates. For instance, a 15 degree vertical bar projected in front of the fly would actually subtend 15.15 degrees because the fly head is a little bit closer than the center of the arena. As the fly rotates, the bar rotates with it and therefore keeps that maximum subtended angle. This system does not fully erase the perception of all visual movement, but it does cause strong disruption to reflexive visual stability demonstrated within the data we present here, as well as those presented previously using this system (48), and a similar approach using an LED based system (29).

### Odor stimuli

A gas multiplexer (Sable Systems) was used to switch mass flow regulated air (Sable Systems MFC-2) through a bubbler of apple cider vinegar or water at the rate of 14 mL/min. The odor port delivering the headspace of the odorant was positioned 3 mm away from the head of the fly. A 5 mm diameter tube below the fly provided a vacuum flow 30 ml/min to clear the odor plume. To measure the spatial distribution of the plume, we used a miniature photoionization detector (mini-PID Aurora Scientific, Ontario, Canada). The detector probe was placed at each point of a 12×12 horizontal grid at the plane of the fly’s antennae. Ethanol, which has an ionization-potential of 10.62 eV, was used as a tracer gas. The odor signal peaks near the fly’s head, but is detectable throughout the full azimuth of the free-yaw fly, which is evident in the sharply directed “anti-tracking” focused 180° from the plume that we have observed in response to aversive benzaldehyde(20).

### Saccade Analysis

Saccade statistics were extracted in multiple phases based on the procedure in (24)and detailed in our freely available custom analysis script written in Python. Angular velocity was calculated as the gradient of the fly headings after applying a 15 Hz Butterworth low-pass filter. Saccades were identified in the heading data by applying a local maximum detector (scipy.signal.find_peaks) to the angular speed. The parameters of this peak-finding function were manually tuned to include as many saccades as possible while excluding large deviations indicative of tracking errors (for 60 fps, distance: 15 frames, width: 2 frames, prominence: [1,30] rads, wlen: 15 frames). This function also provides the left and right bases (adjacent local minima) of the segment containing the peak, offering starting points for measuring the beginning and end of each saccade. Each saccade is processed individually, considering the segment defined by the left base - 10 frames and right bases + 10 frames in order to roughly measure the distribution of velocities before and after the supposed saccade. Using each saccade-specific distribution of pre- and post-saccade angular velocities, we define the beginning and end of the saccade as the duration when angular velocity exceeds 2 standard deviations from the mean. Previous methods, which relied solely on angular speed, were insufficient to identify saccades in the visually-clamped condition because visually-clamped flies often perform saccades superimposed on substantial constant-velocity drifts.

The results were visually-inspected for a random sample of about 10 flies in each of the experiments involving saccade analysis to validate this procedure. The beginning and end time points were used to calculate pertinent parameters to compare saccade statistics across genotypes and visual conditions. For each saccade, we define the following parameters: *starting angle* is the fly’s heading at the start of the saccade, *peak velocity* is the velocity corresponding with the maximum angular speed during the segment after applying a 0-order cubic spline interpolation (scipy.interpolate.interp1d), and *inter-saccade interval (ISI)* is the amount of time since the end of the previous saccade. Given this definition of ISI, the ISI of the first saccade of in each test is undetermined.

### Statistics

All of the statistics presented here amount to comparing means, accounting for within-subject covariances whenever possible. Because many of the variables tested (probability, angular error, normalized cross-correlation, tracking performance index, saccade frequency, peak velocity, and ISI) follow largely non-normal distributions due to mathematical and physical boundaries, we used a bootstrapping approach (54) to test for statistical significance. To do this, the distributions of mean values were sampled with replacement in sets of size N for a total of 100,000 times where N is the sample size. Taking the average of each resampled set generates an empirical sampling distribution of the mean with 100,000 estimates. We then estimate the 95% confidence interval by taking the 2.5 and 97.5 percentiles of the sampling distribution as the lower and upper bounds. Whenever we compare two distributions of mean values, if drawn from independent samples, we compare the sampling distributions elementwise and then calculate the null probability.

For instance, if we hypothesize that some mean value is greater for group A than B our null hypothesis is that 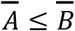. To test this, we would calculate each sampling distribution independently and then store a 1 if A_i_ ≤ B_i_ and 0 otherwise, where A_i_ and B_i_ represent the i-th element in each array. Taking the pseudocount-corrected average provides the null probability:

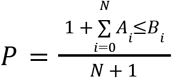

Note that simply taking the average could incorrectly suggest that a result is impossible rather than just not observed in our sample(55).

When comparing results within subjects, we take the within-subject differences first and then calculate P in much the same way, except that our null hypothesis is a comparison to 0 (for instance, *Ā* ≤ 0). When comparing variables with many values per subject—such as the histograms in Figures 2-5 or the time series in Figure 6—sampling is performed hierarchically to maintain the covariances in the resultant sampling distribution. For instance, a histogram was measured for each of N subjects from two different epochs (0-5 and 15-20 s). This resulted in two sets of 11 probabilities, which were randomly sampled in their entirety 100,000 times, resulting in a matrix of size 11 x 2 x N x 100,000. To get the sampling distributions for each mean at each position and epoch, we take the mean along the 3rd dimension, reducing the size of the matrix to 11 x 2 x 100,000. Finally, we get the 95% confidence intervals by taking the 2.5 and 97.5 percentiles along the last dimension, resulting in 2 matrices of size 11 x 2. For histogram comparisons, we actually use 84% confidence intervals because separation of two such intervals indicates a significance level of .05, whereas 95% confidence intervals would represent an overly conservative estimate(56).

## Acknowledgments

This work is funded by grants from the National Institutes of Health R01EY026031 to M.A.F., US Air Force Office of Scientific Research FA9550-23-1-0401 to M.A.F., and National Science Foundation 2016188 to S.M.W. We thank Sarah Fatkin and Martha Rimniceanu for preliminary results.

## Notes

### Competing Interest Statement

The authors have declared no competing interest.

